# Forbs in Viking lands: The effect of disturbing dominant graminoids on forb recruitment in tundra grasslands

**DOI:** 10.1101/2024.12.17.628929

**Authors:** Gerardo Celis, Dorothee Ehrich, Mary Heskel, Eeva Soininen, Peter Ungar, Kari Anne Bråthen

## Abstract

Certain graminoids can be successful in grasslands to the extent it is a phenomenon called “the Viking Syndrome”. Nevertheless, forbs also make up a substantial part of vascular plant diversity in grasslands and are important resources of mammalian herbivores. Here we assess the hypothesis that forb recruitment is constrained by dominant graminoids, limiting access to safe sites for germination. We report on a disturbance experiment of plots with four different graminoid species in tundra grasslands of the Varanger Peninsula, Norway. Plots were selected to sample both rodent-disturbed and undisturbed areas. The dominant graminoids in each plot were removed, reducing their shading capabilities and belowground rhizome and root systems. Results show that forb recruitment one year following disturbance was significantly enhanced by manual graminoid removal. Dominant graminoid type, small rodent disturbance, initial forb abundance, and abiotic conditions had no effect on forb recruitment, whereas initial species richness had a positive relationship. Furthermore we found that manual disturbance had low impact on the species exchange ratio based on richness (SERr), suggesting that disturbance did not reduce the capacity of species to reside and move within the grassland. Our findings support the hypothesis that forb recruitment is limited by dominant graminoids.

## Introduction

Forbs make up a substantial part of the diversity of grasslands, yet they are often low in abundance compared to other functional plant types (Seabloom et al. 2013, Bråthen et al. 2021). Some forb species have such low abundance in grasslands that they are red listed (Eriksson 2013). Nevertheless, the high nutrient content and diversity of forbs suggests they are important to biodiversity and ecosystem functioning, especially in regard to herbivory (e.g., Bråthen et al 2021). And forbs have been found to increase their fitness under disturbance by herbivores (e.g., Inouye 1982, Lennartson et al 1998), emphasizing the role of plant herbivore interactions in the evolution of forbs. It has even been speculated that forbs evolved under conditions of megafaunal disturbance during the Pleistocene, and that their low abundance today may be explained by the extinction of those large herbivores (Bråthen et al. 2021). Addressing the causes of the low abundance of forbs in contemporary grasslands is thus urgent both in terms of knowledge for preventing loss of species and for potential reestablishment of biodiversity and ecosystem functioning in the face of climate change.

Regeneration conditions (Olff and Ritchie 1998, Turnbull et al. 2000) and access to light (Borer et al. 2014) are linked to higher species richness of forbs in grasslands today. Regeneration requires safe germination sites or gaps (Olff and Ritchie 1998, Turnbull et al. 2000), including access to belowground resources (Bråthen et al 2021). However, dominant plant species can limit access to safe germination sites through both shading and belowground root and rhizome networks, preventing less competitive species access to the habitat. Indeed, disturbance of dominant plant species by large herbivores (adult body mass >45 kg) have promoted plant species richness in grasslands (Koerner et al. 2018), including forbs (Bråthen et al. 2021). Here we evaluate this hypothesis with an *in vivo* experiment simulating disturbance of arctic tundra grass plots by megaherbivores to determine the extent to which disruption of dominant plant species limits seedling recruitment of forbs.

Graminoids often colonize, modify, and dominate tundra settings, a phenomenon referred to as “the Viking Syndrome” (Linder et al. 2018). Dominant graminoids often take a rhizomatous (extravaginal tillering) or caespitose (intravaginal tillering) growth form (Klimešová et al. 2017). Clonal rhizomatous graminoids make dense mats often occupying upper soil horizons, giving them a competitive advantage and the capacity to suppress plant species richness (Eilts et al. 2011). Clonal caespitose graminoids, or bunch/tufted graminoids, make dense turfs to the extent that they can cause transformation of the environment (Linder et al. 2018). Depending on the height of the graminoids, they are also potentially strong competitors for light. Furthermore, several dominant graminoids have silica-rich leaves, causing their palatability to be low for herbivores for most of the growing season (Hodson et al. 2005). This suggests that dominant graminoids can have yet another advantage compared to more palatable forbs, underscoring the role of graminoids in potentially limiting forbs in grasslands.

In this study we assess the role of dominant graminoids in inhibiting forb recruitment in tundra grasslands. Forbs are the most nutritious growth form in the tundra and, unsurprisingly, are preferred plant types for both small and large mammalian herbivores (Soininen et al. 2013, Bernes et al. 2015, Bråthen et al. 2017, Petit Bon et al. 2020, Post et al. 2021). This highlights the essential role of these plants for ecosystem functioning, even in tundra grasslands. For instance, despite the dominance of graminoids in their ranges, tundra voles (*Microtus oeconomus*) selectively feed on less abundant forbs, followed by grasses and then deciduous shrubs, with little variation in seasonal (autumn – summer) or spatial preference (river catchments) (Soininen et al. 2013). Forbs are also affected by arctic greening with the encroachment of shrubs limiting the availability of their habitats (Walker et al. 2006, Elmendorf et al. 2012, Berner et al. 2020). Nevertheless, herbivores can hold the shrubs in a browse trap and keep tundra grasslands open (Ravolainen et al. 2014, Bernes et al. 2015, Bråthen et al. 2017), though they can also avoid and hence facilitate the dominance of silica-rich graminoids (Ravolainen et al. 2011). Small rodents in particular have been shown to enhance nutrient turnover and increase the relative abundance of forbs in low productive tundra heathlands through disturbance and feces deposition (Tuomi et al. 2019). They disturb tussock forming graminoids by eating them, but also more importantly by generating litter for shelters and creating burrows and runways through the tussocks (Gough et al. 2007, Lindén et al. 2021). While interactions between plants and herbivores and between plant functional groups in tundra grasslands are complex, there is value in isolating effects of individual interactions on forbs in particular, given their importance to herbivores and changing conditions in low arctic habitats today. Here we aim to isolate the effect of dominant graminoids on limiting forb recruitment by through an experiment involving manual disturbance beside naturally occurring disturbance by small rodents.

We established an experiment in tundra grasslands of Northern Fennoscandia, manually creating gaps or safe sites for seedling establishment. We further evaluated the extent to which forb recruitment in safe sites is dependent on the type of dominant graminoid growth form (mat versus bunch), silica content (rich versus poor) and nutrient enhancement from small rodent disturbance (Petit Bon et al. 2020). In addition, we assessed the impacts of abiotic conditions (light and moisture) on the recruitment process. We predicted that after manual removal of a dominant graminoid, forb recruitment would be a) independent of graminoid growth form, b) independent of graminoid silica content, and c) enhanced by small rodent disturbance. We furthermore predicted that the light intensity and moisture level of the habitat would impact forb recruitment. Besides forb recruitment, we assessed the effect of treatments on the richness- and abundance-based species exchange ratio (SERr and SERa respectively), expecting both the manual and small rodent disturbance to enhance both exchange ratios.

## Material and Methods

### Study site

This study was conducted in riparian grasslands along tributaries of the Bergebyelva river, Varanger Peninsula, Norway (70.30° N, 29.02 ° E, Fig 1). These grasslands harbor dominant graminoids such as *Deschampsia cespitosa, Carex nigra, Calamagrostis spp. and Anthoxanthum nipponicum* and a diverse communityof other plant species, including forbs (e.g.*, Rumex acetosa, Trollius europaeus, Viola* spp.), cryptograms (e.g., *Equisetum* spp. and mosses) and other graminoids (e.g., *Avenella flexuosa, Phleum alpinum, Carex spp. Juncus filiformis)*. The bedrock is a mixture of rich slate and limestone dominated types and poor sandstone types (Ims et al. 2013), see supporting information for climatological summaries of temperature and precipitation. These valleys are part of the study area of the Climate-ecological Observatory for Arctic Tundra (COAT, www.coat.no), which is a long-term ecosystem-based monitoring system addressing Arctic food webs under climate change. The study was initiated in 2023, a peak year of the cyclic small rodent populations in the region (https://coat.no/en/Small-rodent). These tundra grasslands are a primary habitat of the tundra vole, but the Norwegian lemming (*Lemmus lemmus*) and grey-sided vole (*Myodes rufocanus*) are also present. Other vertebrate herbivores found in the study area include semi-domesticated reindeer (*Rangifer tarandus*), moose (*Alces alces*), willow ptarmigan (*Lagopus lagopus*), rock ptarmigan (*Lagopus muta*), and hare (*Lepus timidus*).

**Figure 1.**
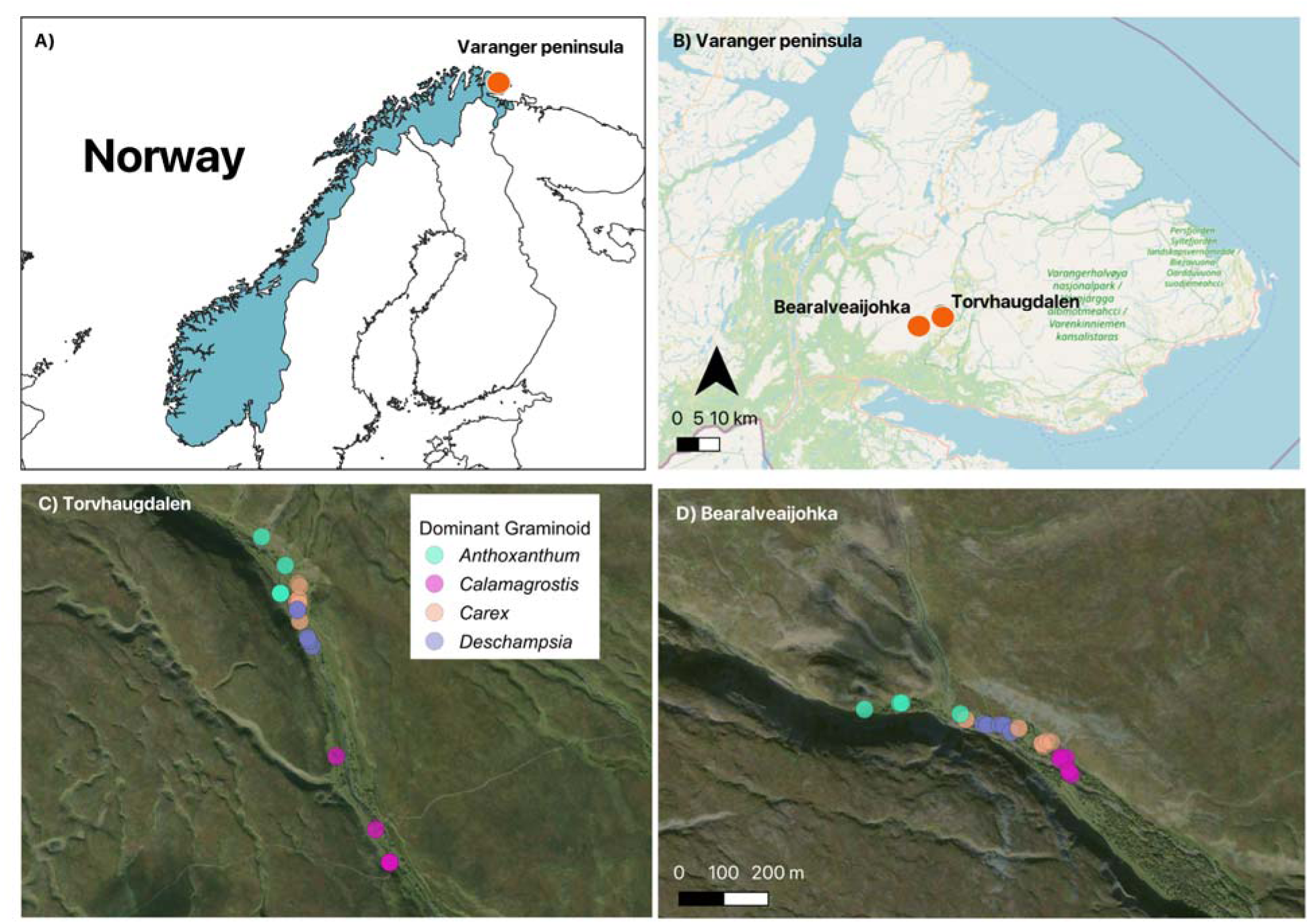
Location of plots in riparian grasslands along Bearalveaijohka and Torvhaugdalen, two tributaries of the Bergebyelva river, Varanger Peninsula, Norway (70.30° N, 29.02 ° E). Source (OpenStreet Maps panel B and ESRI World Imagery panels C-D).

### Experimental design

The main treatment was manual disturbance of dominant graminoids making up most of the vascular plant abundance in grassland vegetation plots. Both aboveground and belowground plant parts were removed, and in doing so most other vascular plants present in the plots were also disturbed (see supporting information with pictures from treated and untreated plots). To test theextents to which small rodent disturbance, dominant graminoid growth form and dominant graminoid silica content affected the outcome of this manual disturbance, we established a nested study design in two riparian valleys (Torvhaugdalen, 70.31° N, 29.08° E and Bearalveaijohka 70.29° N, 28.96° E) in July 2023 (Figure 2, see supporting information). In both valleys, we identified substantial populations of the four dominant graminoids: the silica-rich bunch graminoid *Deschampsia cespitosa*, the silica-poor bunch graminoid *Carex nigra*, the silica-rich mat graminoid *Calamagrostis* spp. and the silica-poor mat graminoid *Anthoxanthum nipponicum*. For each dominant graminoid within each site, we identified areas with no visual signs of rodent disturbance and areas with high levels of rodent disturbance (hereafter denoted as *Non-rodent* and *Rodent* Disturbed). Disturbance was identified by vegetation clipping, runways, and feces presence. In each area we established a minimum of two 0.5m x 0.5m plots, or plot pairs. The plots representing a single pair were at least 0.5m from each other and plot pairs that were either Non-rodent or Rodent Disturbed were at least 7m from each other.

**Figure 2.**
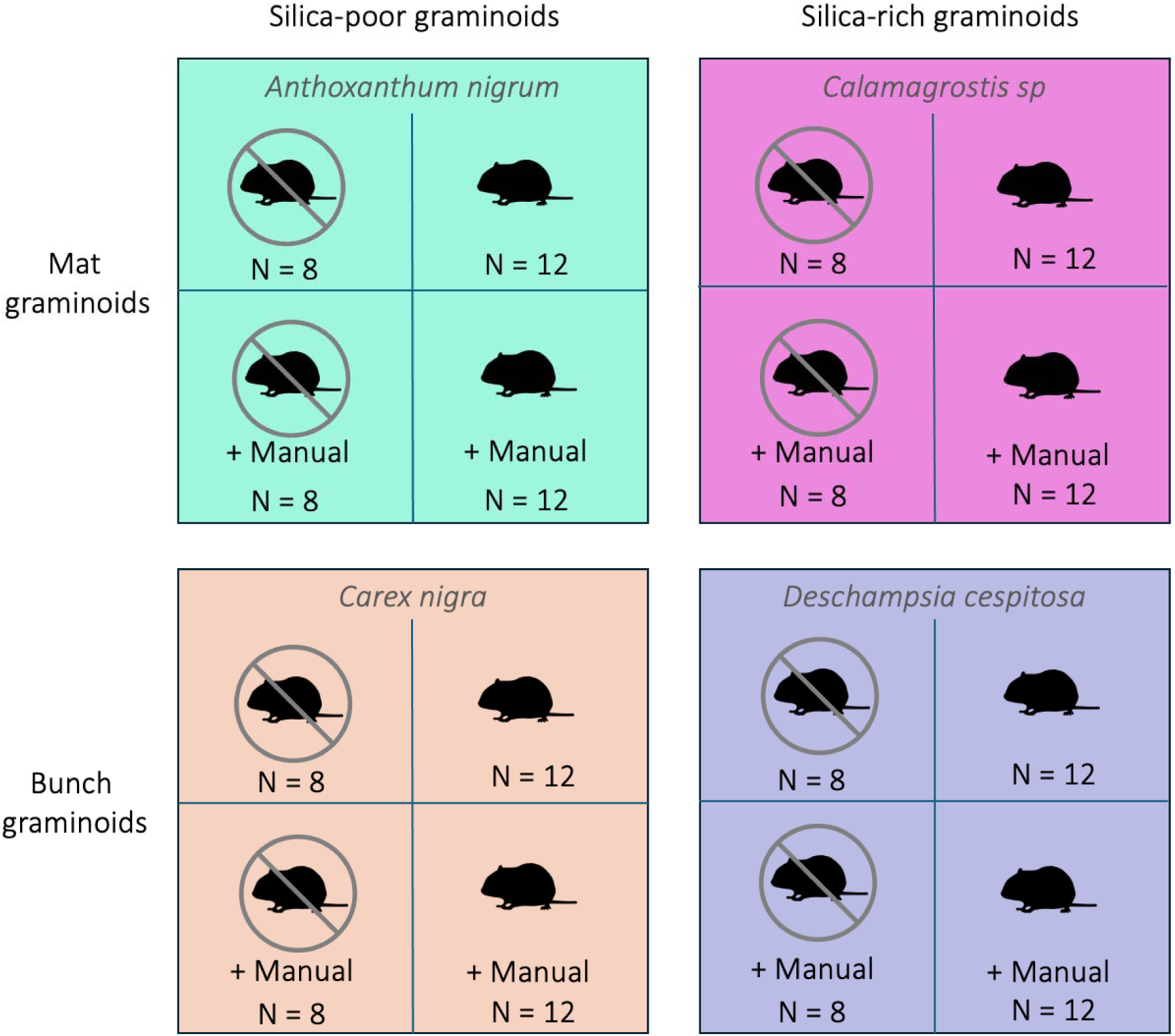
Experimental design for combination of graminoid form, silica content, and rodent disturbance. Small rodent silhouette is from PhyloPic (https://www.phylopic.org/; T. Michael Keesey, 2023) made by Edwin Price, 2023 (CC0 1.0). Treatment colors match those of Figure 1.

Initial plot conditions in July 2023 were determined by recording a list of all vascular plant species as a measure of species richness, estimating the percent cover per species, measuring plant height (in each plot quadrant), and assessing rodent disturbance intensity–percentage of the plot with graminoid clippings or runways and feces presence. After these initial recordings, we randomly chose one plot per pair for the manual disturbance. To simulate megaherbivory capable of disturbing both above- and belowground plant structures, we removed both above- and belowground biomass – breaking shade aboveground and rhizome and belowground root and rhizome networks (Fig. S3 & S4) (hereafter denoted *Manual*).

In July 2024, we revisited each plot, assessed percent cover per species and plant height as in 2023, and counted the number of seedlings in the plot (many seedlings had developed their first true leaf, allowing us to distinguish between forbs and other Dicotyledoneae plants - e.g., *Salix* shrubs). We did not register graminoid seedlings. This was because we could not differentiate clearly between true seedlings and regeneration from buds. Indeed, there were likely remains of rhizomes in the soil after the manual disturbance that have likely survived the winter.

We also sampled plot level abiotic information. We measured photosynthetic active radiation (PAR) at seedling height and above the vegetation canopy using the quantum sensor on a LiCOR 600 Porometer/Fluorometer (LiCOR Biosciences, Lincoln, Nebraska, USA) to determine the relative amount of light reaching seedlings. The PAR index ranged from 0 to 1, 1 representing a situation where all light reaches seedling. We further measured percent soil moisture using a SM150 soil moisture sensor (Delta-T Devices, Cambridge, UK), taking the mean of three measures per plot.

### Species composition, abundance, and turnover

We calculated species richness and Simpson’s (1949) diversity index for each plot for initial conditions in 2023 before any manual treatment. We then estimated indices for species turnover in relation to species identity and dominance shifts, considering initial plot conditions in 2023 and change observed in 2024. We used the species exchange ratio based on richness (SER_r_) and abundance (SER_a_) per plot (Hillebrand et al. 2018) to evaluate species turnover per plot.

### Statistical analyses

Generalized linear mixed models were used to evaluate the effects of dominant graminoid form (mat or bunch), silica content (silica rich or silica poor) and disturbance type (rodent or manual) on species richness, diversity (initial conditions in 2023), species exchange ratios (SER_r_ and SER_a_; difference between 2023 and 2024), and number of forb seedlings in 2024. For each response variable the three design contrasts and their interactions were included as fixed effects, and valley identity was included as a random effect. The forb seedling model also included 2024 Soil moisture and PAR ratio, and 2023 species richness as predictors. The *glmmTMB* function of the *glmmTMB* package (Brooks et al. 2024, v1.1.10) using R statistical software (Team 2024v4.4.2) for models and adjusted to specific probability distributions for each response variables, such that richness and diversity used *gaussian*, SERr and SERa a *beta_family*, forb recruitment a *tweedie*. Models were assessed using the simulation-based package *DHARMa* (Hartig 2024v0.4.7) and *tab_model* function of the *sjPlot* (Lüdecke 2024v2.8.16) package for model summaries. The *ggplot2* package (Wickham et al. 2024v3.5.1) was used for figures. For significant treatment effects, we performed a post-hoc contrast pairwise comparison with Tukey adjustment using *emmeans* (*emmeans* package; Lenth 2024v1.10.5) for dominant graminoid disturbance treatment, interaction between silica content and disturbance treatment, and interaction between graminoid form and disturbance treatment.

## Results

### Initial conditions

Initial plot conditions in 2023 showed that the dominant graminoids made up a substantial cover in plots not disturbed by small rodents, but with the mat graminoid silica-poor *Anthoxanthum nipponicum* having a less dominant role compared to the other dominant graminoids (Table 1). Furthermore, *Anthoxanthum* plots were in the upper range of forb cover and species richness and in the lower range of canopy height and species diversity compared to plots of the other dominant graminoids (Table 1). A total of 19 species or 41% of the total, were shared between graminoids types, and *Anthoxanthum* had 9 unique species, *Carex* 4, *Calamagrostis* 1, and *Deschampsia* did not have any unique species. Within each graminoid type, the cover of forbs and the species richness were similar among the non-rodent and rodent plots (Table 1).

**Table 1.** Initial plot conditions in 2023 before *Manual* disturbance treatment was applied. Values are means (95% confidence interval) for dominant graminoid percent cover, forb percent cover, percent of herbivory, canopy height, species richness and Simpson’s diversity index.

| Dominant graminoid | Disturbance | Dominant cover (%) | Forb cover (%) | Herbivory (%) | Canopy height (cm) | Species richness | Species diversity |
| --- | --- | --- | --- | --- | --- | --- | --- |
| *Anthoxanthum nipponicum* | Non-rodent | 33.7<br>(19.8 - 47.7) | 29.9<br>(16.6 - 43.2) | 3.7<br>(1.8 - 5.7) | 17.0<br>(13.1 - 20.9) | 10.9 (9.5-12.4) | 0.44<br>(0.29 - 0.60) |
|  | Rodent | 16.3<br>(9.4 - 23.3) | 31.5<br>(25.3 - 37.7) | 47.9<br>(37.5 -58.4) | 9.2<br>(5.9 - 12.5) | 10.9<br>(8.9 - 12.9) | 0.40<br>(0.28 - 0.52) |
| *Calamagrostis spp.* | Non-rodent | 60.6<br>(51.0 - 70.2) | 36.0<br>(22.6 - 49.4) | 3.8<br>(1.8 - 5.6) | 45.6<br>(37.0 - 54.1) | 6.0<br>(5.2 - 6.8) | 0.42<br>(0.26 - 0.57) |
|  | Rodent | 47.9<br>(33.4 - 62.4) | 24.8<br>(16.9 - 32.7) | 34.1<br>(27.6 - 40.8) | 42.3<br>(34.5-50.0) | 7.3<br>(6.2 - 8.4) | 0.38<br>(0.28 - 0.48) |
| *Carex nigra* | Non-rodent | 78.1<br>(66.7 - 89.5) | 17.4<br>(16.9 - 32.7) | 6.25<br>(0-15.9) | 40.4<br>(34.6 - 55.7) | 8.6<br>(7.0 - 10.2) | 0.77<br>(0.73 - 0.81) |
|  | Rodent | 74.2<br>(60.0 - 88.4) | 18.2<br>(9.8 - 26.6) | 95.8<br>(78.1 - 99.0) | 46.0<br>(36.4 - 55.7) | 7.5<br>(6.4 - 8.6) | 0.49<br>(0.36 - 0.62) |
| *Deschampsia cespitosa* | Non-rodent | 74.4<br>(67.1 - 81.6) | 11.9<br>(8.3 - 15.5) | 6.25<br>(0.7 - 11.8) | 54.6<br>(45.0 - 64.1) | 8.0<br>(6.3 - 9.7) | 0.73<br>(0.62 - 0.84) |
|  | Rodent | 65.0<br>(55.0 - 75.0) | 19.9<br>(11.5 - 28.3) | 95.8<br>(89.7 - 100) | 52.4<br>(41.3 - 61.6) | 8.2<br>(7.6 - 8.8) | 0.62<br>(0.52 - 0.72) |

### Forb recruitment

The manual disturbance treatment caused a significant increase in forb recruitment in both the *Non-rodent + Manual* and *Rodent + Manual* treatments (Table 2, Fig. 3). The plots identified by the silica-poor mat graminoid *Anthoxanthum* had the least forb recruitment (Fig. 3), with an average of 34 seedlings compared to an average above 85 seedlings in plots identified by the other graminoids. Forb recruitment patterns were maintained between disturbance treatments and graminoid growth form (Table 2, Fig. 3). However, there was a positive interaction between the *Rodent + Manual* treatment and graminoid silica content, with plots identified by the silica-rich graminoids having higher forb recruitment with this treatment with an average of 168 seedlings compared to those identified by silica-poor graminoids with an average of 78 seedlings (Table 2, Fig. 3). Moreover, there was a positive effect of species richness at the start of the experiment (2023) on forb recruitment (Table 2, Figure 4), but no relationship with initial forb cover.

**Figure 3.**
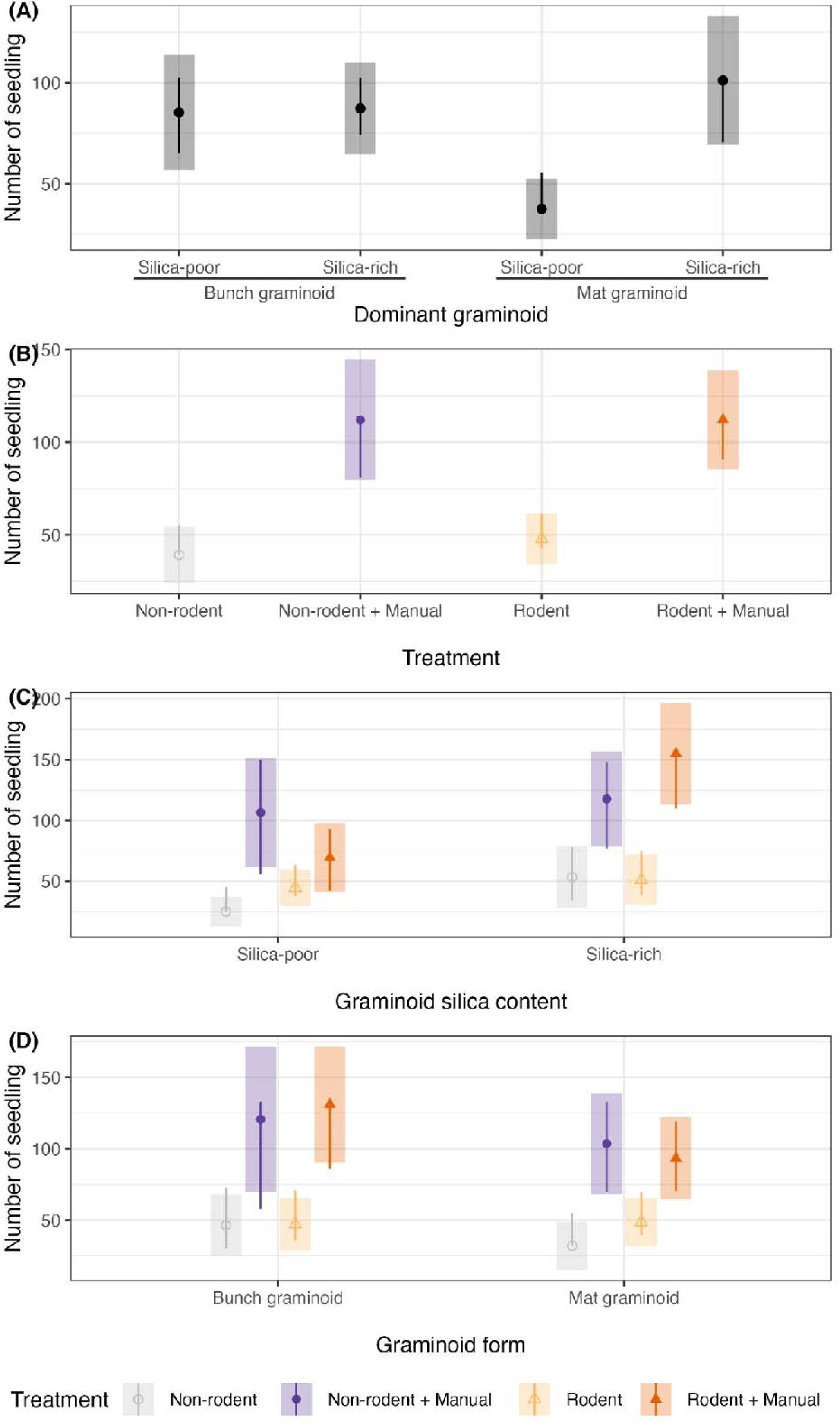
Forb seedling recruitment estimated marginal means based on generalized linear mixed model for each pairwise contrast regarding (A) dominant graminoid; silica-rich bunch graminoid *Deschampsia cespitosa*, silica-poor bunch graminoid *Carex nigra*, the silica-rich mat graminoid *Calamagrostis* spp. and the silica-poor mat graminoid *Anthoxanthum nipponicum*., (B) disturbance treatment (*Non-rodent*, *Non-rodent + Manual*, *Rodent*, *Rodent + Manual*), (C) interaction between silica content (silica-poor and silica-rich) and disturbance treatment, and (D) interaction between graminoid form (mat and bunch) and disturbance treatment. Shaded bars correspond to the 95% confidence intervals for estimates and lines are comparisons among groups, such that if one group line overlaps with another group, the difference is not significant at alpha 0.05, adjusted to Tukey. See supporting information for model details.

**Figure 4.**
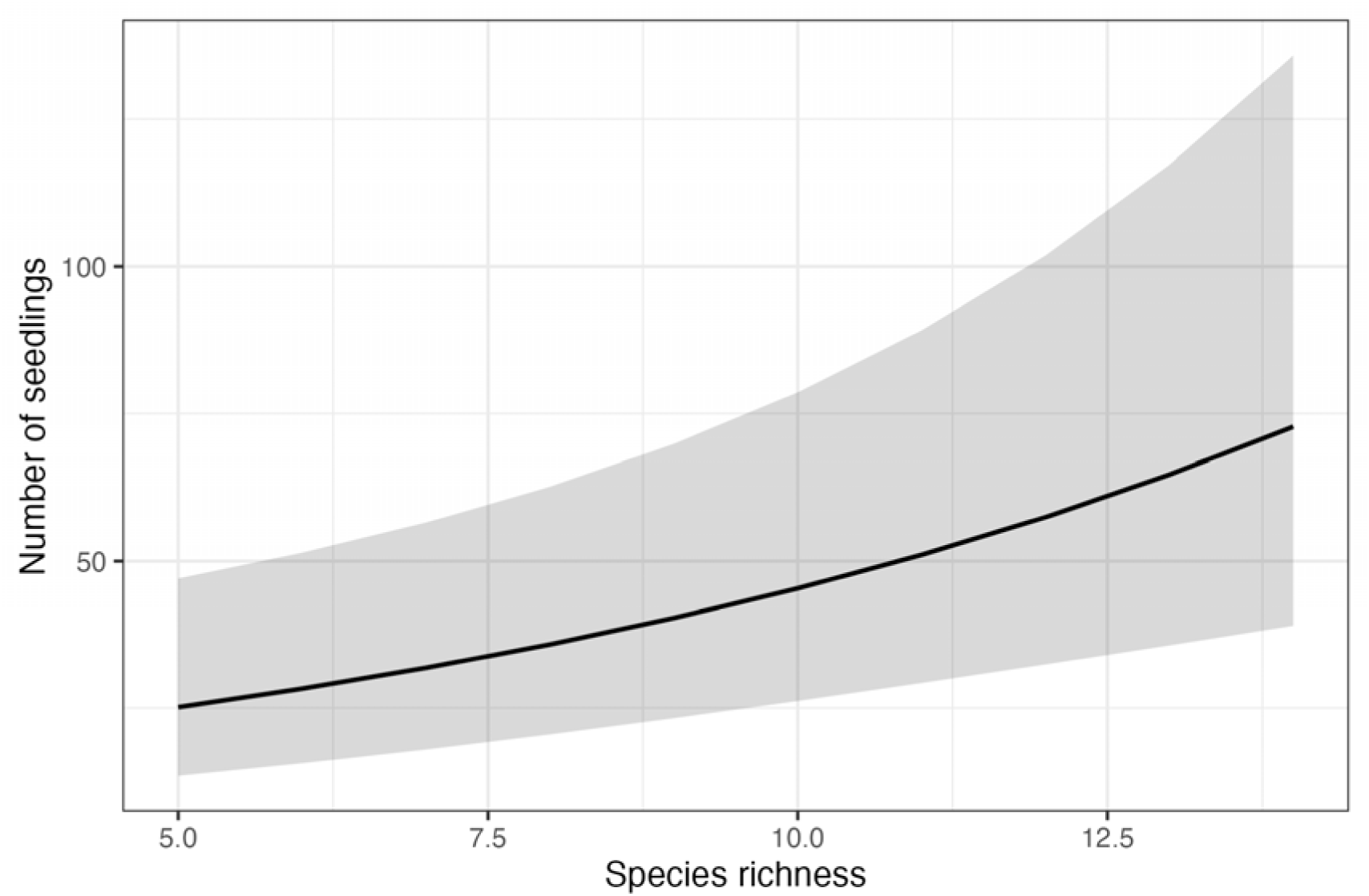
Generalized linear mixed model predicted number of forb seedlings in relation to species richness. See supporting information for model details.

**Table 2.** Generalized linear mixed model results for models evaluating the response of forb seedling recruitment to disturbance treatments, graminoid form, and silica content and their interactions, 2023 forb percent cover, soil moisture (%), photosynthetic active radiation (PAR) ratio, and species richness. Model predictor estimates (on the tweedie scale with link log), confidence intervals (CI) and p-values (p) are reported.

| *Predictors* | **Forb seedling count** |  |  |
| --- | --- | --- | --- |
|  | *Estimates* | *CI* | *p* |
| (Intercept) | 9.14 | 3.91 – 21.39 | **<0.001** |
| TRT [Non-rodent + Manual] | 3.80 | 1.83 – 7.86 | **<0.001** |
| TRT [Rodent] | 1.49 | 0.82 – 2.69 | 0.189 |
| TRT [Rodent + Manual] | 2.76 | 1.36 – 5.60 | **0.005** |
| trt gram [Mat graminoid] | 0.32 | 0.15 – 0.69 | **0.004** |
| trt silica [Silica-rich] | 1.47 | 0.79 – 2.72 | 0.224 |
| Forb percent cover | 1.01 | 1.00 – 1.02 | 0.109 |
| Soil Moisture percent | 1.00 | 1.00 – 1.01 | 0.347 |
| Species richness | 1.13 | 1.05 – 1.20 | **0.001** |
| PAR percent | 1.29 | 0.60 – 2.77 | 0.513 |
| TRT [Non-rodent + Manual] × trt gram [Mat graminoid] | 1.50 | 0.72 – 3.14 | 0.281 |
| TRT [Rodent] × trt gram [Mat graminoid] | 1.81 | 0.87 – 3.75 | 0.112 |
| TRT [Rodent + Manual] × trt gram [Mat graminoid] | 1.03 | 0.50 – 2.10 | 0.936 |
| TRT [Non-rodent + Manual] × trt silica [Silica-rich] | 0.47 | 0.23 – 0.94 | **0.033** |
| TRT [Rodent] × trt silica [Silica-rich] | 0.46 | 0.22 – 0.96 | **0.038** |
| TRT [Rodent + Manual] × trt silica [Silica-rich] | 1.03 | 0.51 – 2.08 | 0.926 |
| trt gram [Mat graminoid] × trt silica [Silica-rich] | 2.87 | 1.54 – 5.35 | **0.001** |
| **Random Effects** |  |  |  |
| $\sigma^2$ | 0.18 | | |
| $\tau_{00}$ valley | 0.01 | | |
| ICC | 0.07 |  |  |
| $N_{\text{valley}}$ | 2 | | |
| Observations | 80 |  |  |
| Marginal $R^2$ / Conditional $R^2$ | 0.689 / 0.709 | | |

### Abiotic conditions

The *Anthoxanthum* plots had on average a higher PAR ratio and lower soil moisture than the the plots of the other dominant graminoids (see supporting information). However, manual treatment of the bunch graminoids caused the PAR ratio to increase more than threefold, and soil moisture levels evinced an increasing trend (the largest differences were found for *Carex nigra* plots) or no change (see supporting information).

There was no significant relationship between the number of seedlings and either PAR ratio or soil moisture (Table 2).

### Species turnover

Species turnover (SERr, based on plot richness) was not significantly affected by manual disturbance, although suggested trends were in general positive, and did not differ between plot categories defined by the graminoid species, with an overall mean ratio of 0.52 (Fig S2, Table S5). However, small rodent disturbance was associated with low SERr, especially in plots with silica-poor graminoids holding a mean ratio of 0.34 (see supporting information).

Species turnover based on plot abundances (SERa) was highest in plots defined by *Anthoxanthum,* the silica-poor mat graminoid, with an average ratio of 0.76 per plot, followed by plots defined by the bunch graminoids with an average ratio of 0.46 and finally plots defined by *Calamagrostis,* the silica rich mat graminoid, with an average ratio of 0.27 per plot. As expected, the manual disturbance treatment increased species turnover in SERa (mean ratio of 0.74 and 0.67 in *Non-rodent + Manual* and *Rodent + Manual* treatments respectively, compared to a mean ratio of 0.29 and 0.24 in *Non-rodent* and *Rodent* treatments respectively.

## Discussion

Forbs are important components of low arctic grassland biodiversity and a key to understanding plant-herbivore interactions on the tundra. These plants are, however, often in relatively low abundance in grasslands due to the “Viking Syndrome”, wherein graminoids colonize, modify, and come to dominate tundra habitats. These dominant graminoids have various growth forms and herbivory defenses that may enhance their competitive advantage over forbs. In this study, we evaluated the effect of dominant graminoids of different growth forms (mat v.s. bunch) and silica content on limiting forb recruitment by conducting an experiment with manual disturbance to simulate megaherbivore activity, beside naturally occurring disturbance by small rodents. The goal was a better understanding how herbivores help mediate the presence of forbs in grasslands.

### Forb recruitment

We found a clear and positive effect of the manual disturbance, that is removal of dominant graminoids, on forb recruitment. The positive effect was independent of graminoid type, although the removal of the smallest of the graminoids (as measured by its average canopy height) had the least positive effect on forb recruitment. Hence, access to safe sites for germination, independent of what type of graminoid had occupied the site before, seems pivotal to forb recruitment.

Plant soil feedbacks from established forbs and graminoids differ, with forbs growing better in soil primed by graminoids than by forbs (Huang et al. 2024). Hence, forb recruitment may have been enhanced in our plots compared to if we had disturbed plots where forbs had previously been dominating. Yet, we found no negative effect of previous forb cover, although the cover was substantial in several plots. This indicates the forb cover was insufficient to cause negative plant soil feedbacks in our study. Negative plant soil feedbacks can however also have been lowered by species richness, diluting negative soil feedbacks (Thakur et al. 2021). Indeed, we found a clear positive relationship between species richness and forb recruitment. On the other hand, graminoids grow equally well in soils primed by either forbs or graminoids (Huang et al. 2024), suggesting they will also have a benefit at the recruitment stage in terms of dominance. It hence remains to see whether the forbs are able to persist past the initial recruitment phase documented here and how the more long term effects of the dominant graminoid removal develop.

Eliminating above-ground vegetation increased access to light for seedlings. However, higher PAR in itself did not promote forb recruitment. Besides having increased access to light, several of the manually disturbed plots had increased soil moisture content, but soil moisture was not related directly to forb recruitment. Germination studies of Arctic forbs have shown that temperature is a primary driver of germination, with higher temperatures resulting in higher germination rates (Müller et al. 2011). Due to its lower albedo, the bare soil can have had higher temperatures after snowmelt, creating optimal conditions for seeds in the soil bank to germinate (Semenchuk et al. 2016), where the balance between access to light and soil moisture may have caused temperatures to be similar in our treated plots.

The small rodent disturbance did not affect forb recruitment significantly in this study, although our models did show a tendency toward a positive effect. This suggests the activities of the naturally occurring small rodents are not enough to enhance forb recruitment to the extent of the more massive disturbance simulating megaherbivory. However, small rodent peak amplitude vary, and 2023 was comparably low peak (Soininen, Ehrich et al. unpublished data). Peak years of small rodents with higher population numbers than that of 2023 may well have a stronger impact. Given that silica-rich graminoids are providing habitats for the small rodents, their presence is potentially also facilitating the small rodents to thrive in the grasslands and promoting their disturbance activities, highlighting the complex interactions between plants and herbivores and between plant functional groups in tundra grasslands.

The generally high species turnover in the grassland plots between the summer seasons of 2023 and 2024 was indicative of a dynamic plant community. Interestingly, and similar to the forb recruitment, species turnover was not affected by the dominant graminoid type. However, only 41% of the species were shared among the plots identified by the four dominant graminoid species. This suggests the turnover dynamics may be dependent on particular species pools associated with the different graminoids, and that future work might benefit from a focus on the habitat context under which the graminoids are dominating.

## Conclusion

Understanding the ability for nutrient-rich forbs to grow and recruit seedlings remains a concern for the future of tundra grassland communities. Here we find a massive increase in forb seedlings in safe sites previously occupied by dominant graminoids, suggesting that dominant graminoids are severely limiting the recruitment of forbs. Hence, disturbing the dominant graminoids can enhance forb recruitment. For forbs to gain a stable presence in the tundra grasslands depends on whether the disturbance also enhances the growth and reproduction of the forbs.

## Acknowledgments

We thank the team of field assistants who helped collect data in 2023 and 2024, including Berit Gaski, Oliver Paine, Lukas Lock-Scamp, Mary Ossar, and Matt Wattendorf, a teacher supported by PolarSTEAM (US National Science Foundation grant 2221990). This study is a contribution from the Climate-Ecological Observatory for Arctic Tundra (COAT)

