## supplemental information for "Forbs in Viking lands: The effect of disturbing dominant graminoids on forb recruitment in tundra grasslands"

**Celis et al., 2024**

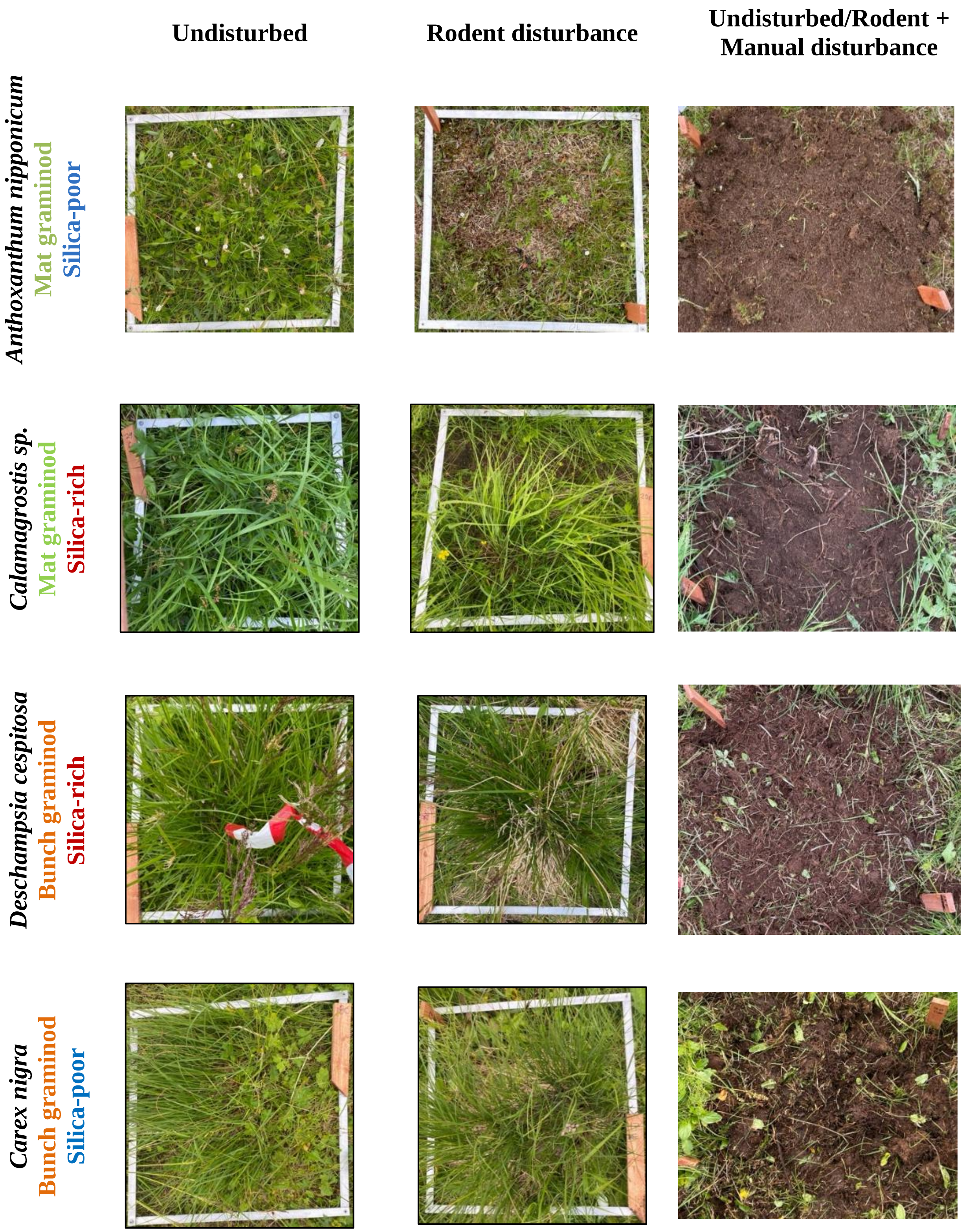


Figure S1. Plot photographs for each dominant graminoid type (rows), and disturbance treatments (columns).

Table S1. The climate normals 1991-2020 period for air temperature and precipitation at study area based on ERA5 data. Values are means and standard errors.

| Period | Temperature (°C) | Precipitation (mm) |
|---|---|---|
| Annual | -1.24 (±0.16) | 633 (±11) |
| Summer (June, July and August) | 8.97 (±0.19) | 212 (±9) |
| July | 10.77 (±0.33) | 73 (±7) |

Table S2. Mean percent cover for each species across all treatments and locations for each year.

| Species | Plant functional group | Species abbreviation | Cover (%) | | Number of Plots | |
|---|---|---|---|---|---|---|
| | | | 2023 | 2024 | 2023 | 2024 |
| *Agrostis sp* | Broad leaved graminoid | agr_sp | 0.0125 | 0.0125 | 1 | 1 |
| *Anthoxanthum nipponicum* | Broad leaved graminoid | ant_nip | 6.0125 | 0.075 | 27 | 2 |
| *Anthoxanthum odoratum* | Broad leaved graminoid | ant_odo | 0 | 0.0875 | 0 | 5 |
| *Calamagrostis sp* | Broad leaved graminoid | cal_sp | 13.4125 | 7.8125 | 22 | 24 |
| *Deschampsia cespitosa* | Broad leaved graminoid | des_ces | 17.7625 | 7.9 | 30 | 22 |
| *Phleum sp* | Broad leaved graminoid | phl_sp | 0.825 | 0.5625 | 19 | 25 |
| *Poa sp* | Broad leaved graminoid | poa_sp | 0.7375 | 0.1 | 18 | 7 |
| *Trisetum spicatum* | Broad leaved graminoid | tri_spi | 0.025 | 0 | 2 | 0 |
| *Avenella flexuosa* | Narrow leaved graminoid | ave_fle | 0.625 | 3.3375 | 4 | 19 |
| *Festuca sp* | Narrow leaved graminoid | fes_sp | 0.0125 | 0.025 | 1 | 1 |
| *Nardus stricta* | Narrow leaved graminoid | nar_str | 0.0125 | 0.425 | 1 | 5 |
| *Carex nigra* | Sedges & Rushes | car_nig | 18.9375 | 9.1125 | 20 | 13 |
| *Carex sp* | Sedges & Rushes | car_sp | 1.1625 | 0.775 | 24 | 19 |
| *Juncus sp* | Sedges & Rushes | jun_sp | 0.1 | 0.1125 | 4 | 3 |
| *Luzula sp* | Sedges & Rushes | luz_sp | 0.1875 | 0.0625 | 10 | 4 |
| *Betula nana* | Deciduous dwarf shrubs | bet_nan | 0.025 | 0 | 2 | 0 |
| *Salix herbaceae* | Deciduous dwarf shrubs | sal_her | 0.2875 | 0.275 | 6 | 8 |
| *Vaccinium myrtillus* | Deciduous dwarf shrubs | vac_myr | 0.825 | 0.275 | 5 | 4 |
| *Vaccinium uliginosum* | Deciduous dwarf shrubs | vac_uli | 0 | 0.0375 | 0 | 2 |
| *Empetrum nigrum* | Evergreen dwarf shrubs | emp_nig | 0.0125 | 0.0125 | 1 | 1 |
| *Vaccinium vitis-idaea* | Evergreen dwarf shrubs | vac_vit | 0.1875 | 0.1 | 2 | 2 |
| *Pyrola sp* | Evergreen dwarf shrubs | pyr_sp | 0.0125 | 0.0375 | 1 | 2 |

| | | | | | | |
|---|---|---|---|---|---|---|
| *Salix sp* | prostrate_willows | sal_sp | 0.325 | 0.1375 | 13 | 5 |
| *Alchemilla sp* | Small forbs | alc_sp | 4.7 | 2.375 | 46 | 39 |
| *Angelica sp* | Small forbs | ang_sp | 0 | 0.0125 | 0 | 1 |
| *Bistorta vivipara* | Small forbs | bis_viv | 2.025 | 0.725 | 38 | 33 |
| *Caltha palustris* | Small forbs | cal_pal | 0.1375 | 0.1625 | 7 | 10 |
| *Cerastium sp* | Small forbs | cer_sp | 0.2875 | 0.025 | 4 | 2 |
| *Chamaepericlymenum suecicum* | Small forbs | cha_sue | 0.625 | 1.0375 | 3 | 4 |
| *Comarum palustre* | Small forbs | com_pal | 0 | 0.05 | 0 | 1 |
| *Epilobium sp* | Small forbs | epi_sp | 0.1375 | 0.5125 | 11 | 27 |
| *Hieracium sp* | Small forbs | hie_sp | 0 | 0.0375 | 0 | 3 |
| *Omalotheca sp* | Small forbs | oma_sp | 0.5625 | 0.375 | 14 | 18 |
| *Ranunculus sp* | Small forbs | ran_sp | 1.3375 | 0.9375 | 34 | 36 |
| *Rubus chamaemorus* | Small forbs | rub_cha | 0 | 0.0125 | 0 | 1 |
| *Rumex acetosa* | Small forbs | rum_ace | 6.3125 | 2.8875 | 64 | 57 |
| *Sagina sp* | Small forbs | sag_sp | 0 | 0.025 | 0 | 2 |
| *Sibbaldia procumbens* | Small forbs | sib_pro | 0.2125 | 0.1125 | 11 | 5 |
| *Solidago virgaurea* | Small forbs | sol_vir | 0.7375 | 0.2375 | 20 | 9 |
| *Stellaria sp* | Small forbs | ste_sp | 0.4875 | 0.7 | 25 | 22 |
| *Taraxacum sp* | Small forbs | tar_sp | 0.7 | 0.5875 | 31 | 26 |
| *Trientalis europaea* | Small forbs | tri_eur | 0.6875 | 0.525 | 20 | 17 |
| *Veronica alpina* | Small forbs | ver_alp | 0.025 | 0.0125 | 2 | 1 |
| *Viola sp* | Small forbs | vio_sp | 0.9125 | 0.675 | 40 | 35 |
| *Viscaria alpina* | Small forbs | vis_alp | 0.0125 | 0 | 1 | 0 |
| *Anthriscus sp* | Tall forbs | ant_sp | 0.125 | 0.875 | 1 | 19 |
| *Chamerion angustifolium* | Tall forbs | cha_ang | 0 | 0.0125 | 0 | 1 |
| *Cirsium heterophyllum* | Tall forbs | cir_het | 0.5875 | 0.675 | 7 | 6 |

| | | | | | | |
|---|---|---|---|---|---|---|
| *Geranium sylvaticum* | Tall forbs | ger_syl | 1.4875 | 0.475 | 19 | 11 |
| *Trollius europaeus* | Tall forbs | tro_eur | 1.5875 | 1.0625 | 15 | 16 |
| *Equisetum sp* | equ_sp | equ_sp | 3.2125 | 3.0125 | 35 | 44 |
| | | Moss | 0.375 | 0 | 1 | 0 |

Table S3. Summary of variables for each disturbance treatment, graminoid form, and silica content. Values are means (95% confidence interval) for forb percent cover and percent of herbivory in 2023, Species exchange ratio based on species richness (SERr) and abundance (SERa) between 2023 and 2024, and soil moisture percent and PAR ratio measured in 2024

| Dominant graminoid | Treatment | 2023<br>Forb cover (%) | 2023<br>Herbivory (%) | 2023 - 2024<br>SERr | SERa | 2024<br>Soil Moisture (%) | PAR ratio |
|---|---|---|---|---|---|---|---|
| Mat, Silica-poor<br>Anthoxanthum nipponnicum | Non-rodent | 35.8<br>(24.3 - 47.3) | 3.8<br>(-0.2 - 7.8) | 0.63<br>(0.58 - 0.68) | 0.86<br>(0.78 - 0.94) | 11.39<br>(10.2 - 12.5) | 0.59<br>(0.38 - 0.8) |
| | Non-rodent + Manual | 24.0<br>(11 - 37) | 3.8<br>(-0.2 - 7.8) | 0.74<br>(0.69 - 0.79) | 0.90<br>(0.86 - 0.94) | 13.2<br>(10.9 - 15.6) | 0.87<br>(0.76 - 0.98) |
| | Rodent | 30.3<br>(23.3 - 37.3) | 47.5<br>(30.7 - 64.3) | 0.41<br>(0.33 - 0.49) | 0.63<br>(0.49 - 0.77) | 9.4<br>(6.6 - 12.2) | 0.87<br>(0.8 - 0.94) |
| | Rodent + Manual | 32.7<br>(27.6 - 37.8) | 48.3<br>(29 - 67.6) | 0.60<br>(0.51 - 0.69) | 0.70<br>(0.6 - 0.8) | 10.38<br>(5.9 - 14.9) | 0.92<br>(0.88 - 0.96) |
| Mat, Silica-rich<br>Calamagrostis sp | Non-rodent | 35.2<br>(22.2 - 48.2) | 3.8<br>(-0.2 - 7.8) | 0.37<br>(0.24 - 0.5) | 0.14<br>(0.05 - 0.23) | 21.5<br>(15 - 28) | 0.15<br>(0.07 - 0.23) |
| | Non-rodent + Manual | 36.8<br>(22.9 - 50.7) | 3.8<br>(-0.2 - 7.8) | 0.47<br>(0.35 - 0.59) | 0.44<br>(0.2 - 0.68) | 33.0<br>(25.5 - 40.5) | 0.31<br>(0.25 - 0.37) |
| | Rodent | 30.7<br>(24.2 - 37.2) | 30<br>(21.2 - 38.8) | 0.46<br>(0.35 - 0.57) | 0.17<br>(0.05 - 0.29) | 23.6<br>(13.5 - 33.6) | 0.20<br>(0.12 - 0.28) |
| | Rodent + Manual | 19.0<br>(11.7 - 26.3) | 38.3<br>(26.5 - 50.1) | 0.58<br>(0.49 - 0.67) | 0.34<br>(0.19 - 0.49) | 28.1<br>(16.8 - 39.4) | 0.31<br>(0.24 - 0.38) |
| Bunch, Silica-poor<br>Carex nigra | Non-rodent | 15.5<br>(9.6 - 21.4) | 6.2<br>(-13.7 - 26.1) | 0.50<br>(0.3 - 0.7) | 0.06<br>(-0.01 - 0.13) | 41.6<br>(24.5 - 58.8) | 0.17<br>(0.03 - 0.31) |

| | | | | | | | |
|---|---|---|---|---|---|---|---|
| | Non-rodent + Manual | 19.2<br>(9.7 - 28.7) | 6.2<br>(-13.7 - 26.1) | 0.56<br>(0.52 - 0.6) | 0.89<br>(0.81 - 0.97) | 72.3<br>(45.2 - 99.4) | 0.72<br>(0.63 - 0.81) |
| | Rodent | 16.5<br>(11 - 22) | 87.5<br>(65.5 - 109.5) | 0.27<br>(0.2 - 0.34) | 0.07<br>(0.03 - 0.11) | 46.3<br>(35.1 - 57.6) | 0.21<br>(0.08 - 0.34) |
| | Rodent + Manual | 20.0<br>(9.7 - 30.3) | 89.6<br>(76.7 - 102.5) | 0.58<br>(0.5 - 0.66) | 0.76<br>(0.64 - 0.88) | 74.3<br>(54.9 - 93.7) | 0.63<br>(0.49 - 0.77) |
| Bunch, Silica-rich<br>Deschampsia cespitosa | Non-rodent | 13.8<br>(10.3 - 17.3) | 6.2<br>(-5.3 - 17.7) | 0.59<br>(0.46 - 0.72) | 0.04<br>(-0.01 - 0.09) | 43.0<br>(20.7 - 65.3) | 0.16<br>(0.09 - 0.23) |
| | Non-rodent + Manual | 10 (7.1 - 12.9) | 6.2<br>(-5.3 - 17.7) | 0.61<br>(0.58 - 0.64) | 0.83<br>(0.73 - 0.93) | 53.9<br>(38.8 - 69.0) | 0.64<br>(0.43 - 0.85) |
| | Rodent | 17.5<br>(10.4 - 24.6) | 100<br>(100 - 100) | 0.50<br>(0.41 - 0.59) | 0.15<br>(0.02 - 0.28) | 28.3<br>(19 - 37.7) | 0.17<br>(0.08 - 0.26) |
| | Rodent + Manual | 22.3<br>(13 - 31.6) | 91.7<br>(78.2 - 105.2) | 0.54<br>(0.43 - 0.65) | 0.86<br>(0.79 - 0.93) | 24.0<br>(17.3 - 30.6) | 0.61<br>(0.53 - 0.69) |

Table S4. Generalized linear mixed model results for models evaluating the response of species richness (N) and Simpson's (D) diversity index to Disturbance treatments, graminoid form, and silica content and their interactions for initial plot conditions in 2023. Model predictor estimates (on gaussian scale), confidence intervals (CI) and p-values (p) are reported.

| | **N** | | | **D** | | |
|---|---|---|---|---|---|---|
| *Predictors* | *Estimates* | *CI* | *p* | *Estimates* | *CI* | *p* |
| (Intercept) | 8.31 | 6.80 – 9.82 | **<0.001** | 0.75 | 0.61 – 0.89 | **<0.001** |
| TRT [Non-rodent + Manual] | 0.62 | -1.31 – 2.56 | 0.527 | 0.00 | -0.18 – 0.19 | 0.983 |
| TRT [Rodent] | -0.37 | -2.14 – 1.39 | 0.678 | -0.25 | -0.42 – -0.08 | **0.004** |
| TRT [Rodent + Manual] | -1.25 | -3.02 – 0.52 | 0.166 | -0.23 | -0.40 – -0.06 | **0.007** |
| trt gram [Mat graminoid] | 2.88 | 1.14 – 4.61 | **0.001** | -0.23 | -0.39 – -0.06 | **0.007** |
| trt silica [Silica-rich] | -1.12 | -2.86 – 0.61 | 0.203 | -0.03 | -0.20 – 0.13 | 0.681 |
| TRT [Non-rodent + Manual]× trt gram [Mat graminoid] | -1.25 | -3.49 – 0.99 | 0.273 | -0.11 | -0.32 – 0.11 | 0.333 |
| TRT [Rodent] × trt gram [Mat graminoid] | 0.42 | -1.63 – 2.46 | 0.689 | 0.17 | -0.02 – 0.37 | 0.082 |
| TRT [Rodent + Manual] × trt gram [Mat graminoid] | 0.67 | -1.38 – 2.71 | 0.522 | 0.03 | -0.16 – 0.23 | 0.755 |
| TRT [Non-rodent + Manual] × trt silica [Silica-rich] | 1.00 | -1.24 – 3.24 | 0.381 | 0.08 | -0.13 – 0.30 | 0.436 |
| TRT [Rodent] × trt silica [Silica-rich] | 1.42 | -0.63 – 3.46 | 0.174 | 0.08 | -0.12 – 0.27 | 0.430 |
| TRT [Rodent + Manual] × trt silica [Silica-rich] | 2.17 | 0.12 – 4.21 | **0.038** | 0.19 | -0.01 – 0.38 | 0.060 |
| trt gram [Mat graminoid] × trt silica [Silica-rich] | -4.25 | -5.66 – -2.84 | **<0.001** | -0.08 | -0.22 – 0.05 | 0.217 |
| **Random Effects** | | | | | | |
| $\sigma^2$ | 2.61 | | | 0.02 | | |
| $\tau_{00\ \text{valley}}$ | 0.15 | | | 0.00 | | |
| ICC | 0.05 | | | 0.01 | | |
| $N_{\text{valley}}$ | 2 | | | 2 | | |
| Observations | 80 | | | 80 | | |
| Marginal $R^2$ / Conditional $R^2$ | 0.488 / 0.516 | | | 0.469 / 0.473 | | |

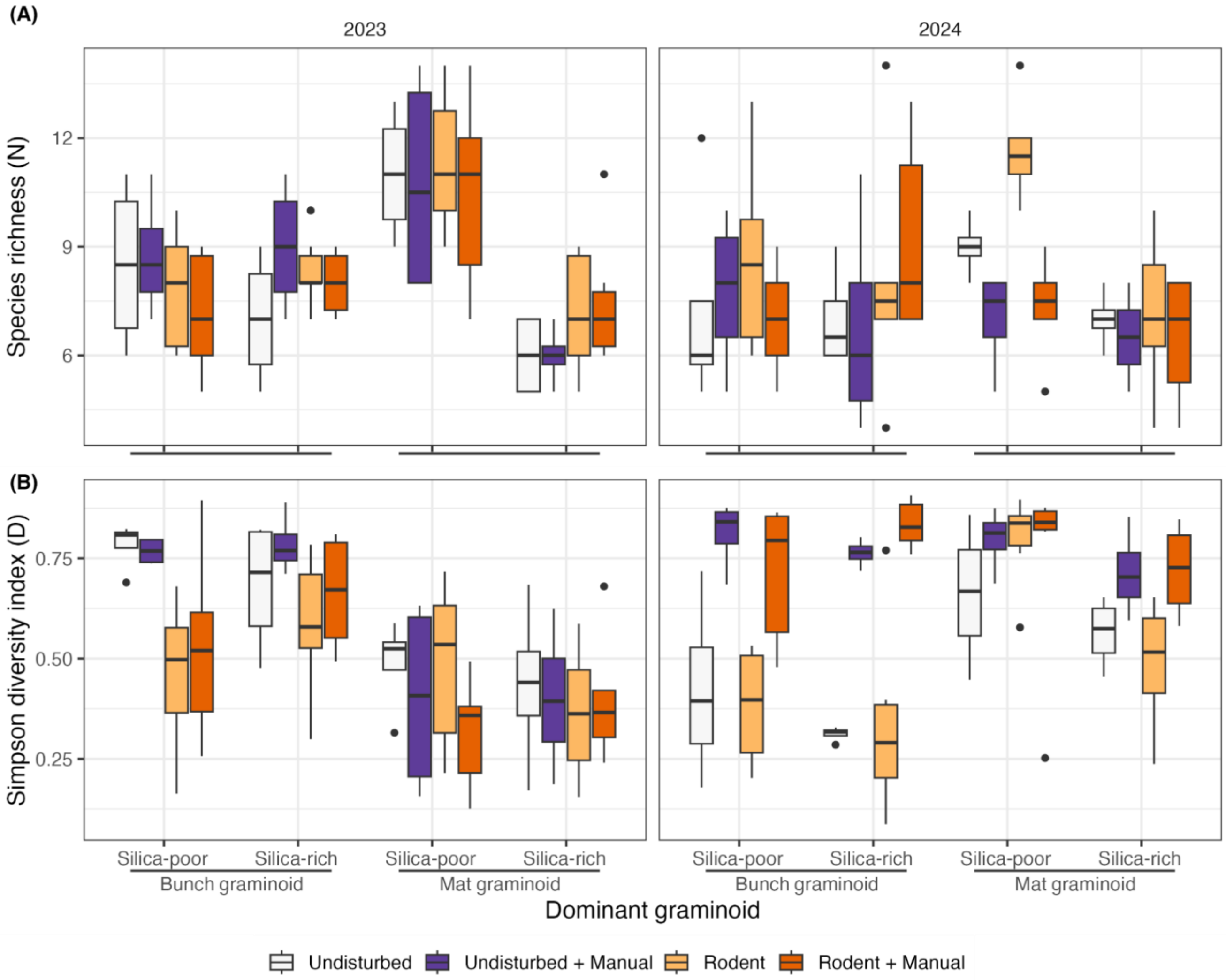


Figure S2. Boxplots (horizontal line is median and box are first and third quartiles) of species richness (A) and Simpson's diversity index (B) per plot (0.5 X 0.5 m) for each dominant graminoid form (mat and bunch), silica content and disturbance treatments (Non-rodent, Non-rodent with manual disturbance, rodent disturbed, and rodent disturbed with manual disturbance) for each year.

Table S5. Generalized linear mixed model results for models evaluating the response of species exchange ratios based on species richness (SERr) and abundance (SERa) to disturbance treatments, graminoid form, and silica content and their interactions for change between 2023 and 2024 plot inventories. Model predictor estimates (on beta scale with link logit), confidence intervals (CI) and p-values (p) are reported.

| | SERr | | | SERa | | |
|---|---|---|---|---|---|---|
| *Predictors* | *Estimates* | *CI* | *p* | *Estimates* | *CI* | *p* |
| (Intercept) | 1.13 | 0.67 – 1.90 | 0.655 | 0.11 | 0.05 – 0.26 | **<0.001** |
| TRT [Non-rodent + Manual] | 1.25 | 0.64 – 2.46 | 0.518 | 50.72 | 16.72 – 153.81 | **<0.001** |
| TRT [Rodent] | 0.34 | 0.18 – 0.64 | **0.001** | 0.91 | 0.33 – 2.48 | 0.853 |
| TRT [Rodent + Manual] | 1.00 | 0.54 – 1.85 | 0.998 | 31.03 | 11.51 – 83.63 | **<0.001** |
| trt gram [Mat graminoid] | 1.19 | 0.65 – 2.17 | 0.567 | 38.07 | 13.97 – 103.78 | **<0.001** |
| trt silica [Silica-rich] | 1.05 | 0.58 – 1.92 | 0.866 | 0.50 | 0.18 – 1.38 | 0.180 |
| TRT [Non-rodent + Manual] × trt gram [Mat graminoid] | 1.34 | 0.62 – 2.92 | 0.462 | 0.03 | 0.01 – 0.11 | **<0.001** |
| TRT [Rodent] × trt gram [Mat graminoid] | 1.50 | 0.74 – 3.07 | 0.263 | 0.43 | 0.13 – 1.38 | 0.158 |
| TRT [Rodent + Manual] × trt gram [Mat graminoid] | 1.37 | 0.67 – 2.79 | 0.385 | 0.02 | 0.01 – 0.06 | **<0.001** |
| TRT [Non-rodent + Manual] × trt silica [Silica-rich] | 0.88 | 0.41 – 1.93 | 0.757 | 1.78 | 0.50 – 6.28 | 0.370 |
| TRT [Rodent] × trt silica [Silica-rich] | 2.48 | 1.22 – 5.07 | **0.012** | 1.99 | 0.63 – 6.34 | 0.242 |
| TRT [Rodent + Manual] × trt silica [Silica-rich] | 1.21 | 0.59 – 2.47 | 0.598 | 3.45 | 1.11 – 10.77 | **0.033** |
| trt gram [Mat graminoid] × trt silica [Silica-rich] | 0.49 | 0.30 – 0.80 | **0.005** | 0.12 | 0.05 – 0.26 | **<0.001** |
| **Random Effects** | | | | | | |
| $\sigma^2$ | 0.07 | | | 0.17 | | |
| $\tau_{00\ valley}$ | 0.02 | | | 0.02 | | |
| ICC | 0.19 | | | 0.13 | | |
| $N_{valley}$ | 2 | | | 2 | | |
| Observations | 80 | | | 80 | | |
| Marginal $R^2$ / Conditional $R^2$ | 0.664 / 0.728 | | | 0.936 / 0.944 | | |

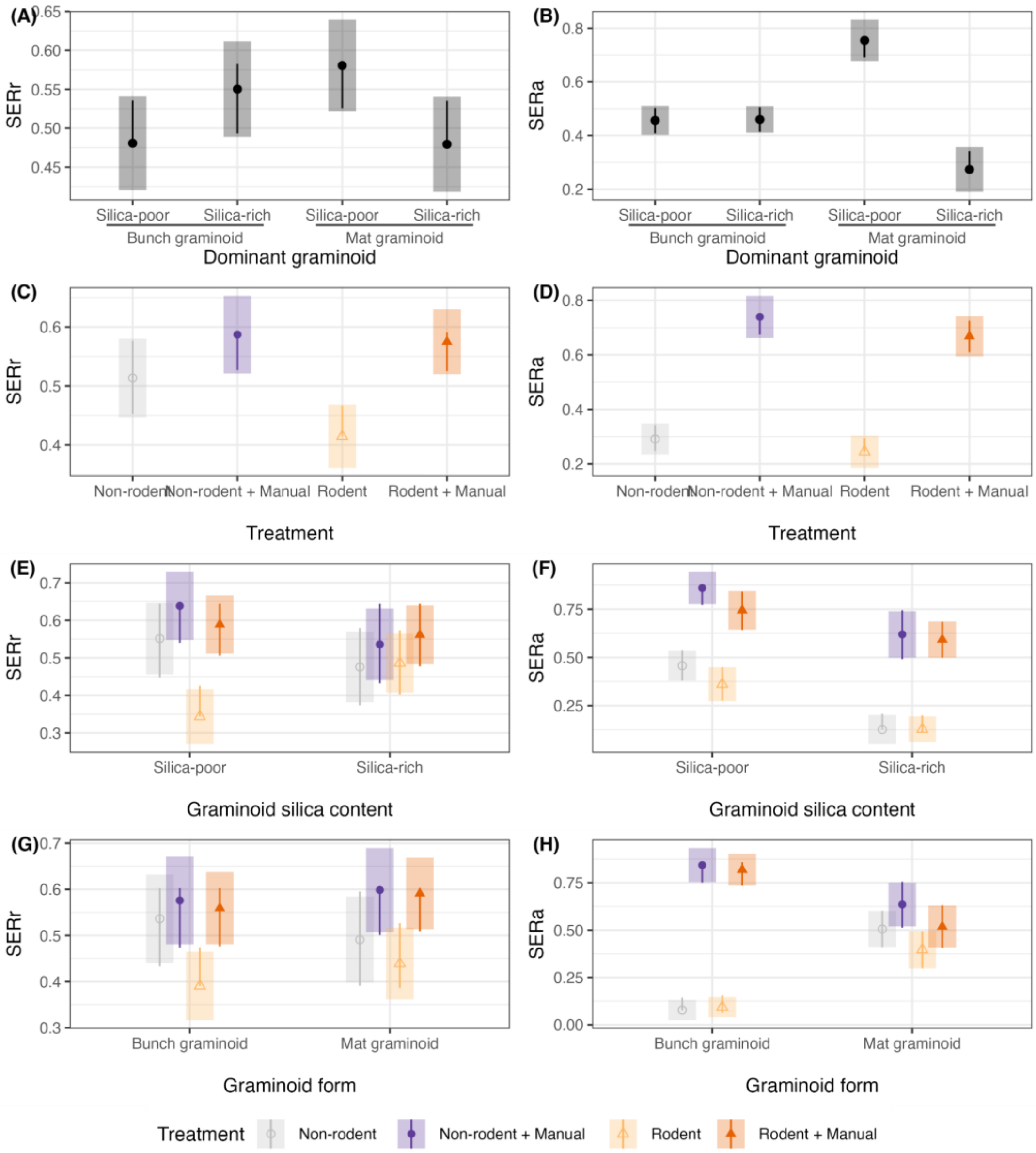


Figure S3. Species exchange ration based on richness SERr and species exchange ration based on abundance SERa estimated marginal means based on generalized linear mixed model for each pairwise contrast regarding (A) dominant graminoid; silica-rich bunch graminoid *Deschampsia cespitosa*, silica-poor bunch graminoid *Carex nigra*, the silica-rich mat graminoid *Calamagrostis* spp. and the silica-poor mat graminoid *Anthoxanthum nipponicum*., (B) disturbance treatment (*Non-rodent*, *Non-rodent + Manual*, *Rodent*, *Rodent + Manual*), (C) interaction between silica content (silica-poor and silica-rich) and disturbance treatment, and (D) interaction between graminoid form (mat and bunch) and disturbance treatment. Shaded bars correspond to the 95% confidence intervals for estimates and lines are comparisons among groups, such that if one group line overlaps with another group the difference is not significant at alpha 0.05 adjusted to Tukey.

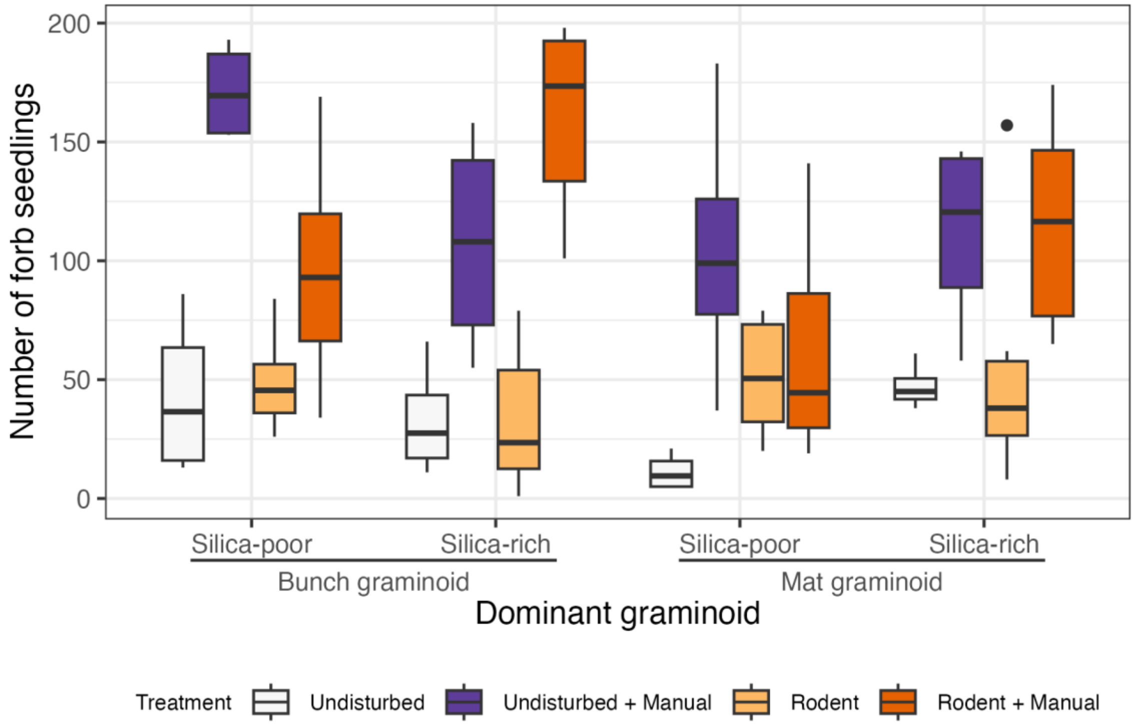


Figure S4. Boxplots (horizontal line is median and box are first and third quartiles) of the number of forb seedling recruitment per plot (0.5 X 0.5 m) for each dominant graminoid form (mat and bunch), silica content and disturbance treatments (*Non-rodent*, *Non-rodent* with *Manual* disturbance, *Rodent* disturbed, and *Rodent* disturbed with *Manual* disturbance).
